# Light-dependent modulation of protein localization and function in living bacteria cells

**DOI:** 10.1101/2022.05.01.490209

**Authors:** Ryan McQuillen, Xinxing Yang, Christopher H. Bohrer, Joshua W. McCausland, Jie Xiao

## Abstract

Most bacteria lack membrane-enclosed organelles to compartmentalize cellular processes. In lieu of physical compartments, bacterial proteins are often recruited to macromolecular scaffolds at specific subcellular locations to carry out their functions. Consequently, the ability to modulate a protein’s subcellular location with high precision and speed bears the potential to manipulate its corresponding cellular functions. Here we demonstrate that the CRY2/CIB1 system from *Arabidopsis thaliana* can be used to rapidly direct proteins to different subcellular locations inside live *E. coli* cells including the nucleoid, the cell pole, membrane, and the midcell division plane. We further show that such light-induced re-localization can be used to rapidly inhibit cytokinesis in actively dividing *E. coli* cells. Finally, we demonstrate that the CRY2/CIBN binding kinetics can be modulated by green light, adding a new dimension of control to the system.

## INTRODUCTION

Compartmentalizing biological processes at specific subcellular locations for specialized functions is a general strategy employed by cells across all domains of life. Eukaryotic cells often achieve subcellular localization using membrane-enclosed organelles. Bacteria cells generally lack membranous organelles, but can also achieve subcellular localization by directing molecules to different scaffolds made of protein, DNA, RNA, or the plasma membrane to form stable assemblies and/or phase-separated condensates^1^. These subcellular scaffolds underlie a number of fundamental cellular processes including partitioning of cellular content, chromosome organization and segregation, cell elongation and division, as well as appendage growth and receptor clustering^1,2^. Controlling the recruitment of proteins to subcellular scaffolds and/or modulating scaffolds’ assembly states has proven to be a useful tool to probe and manipulate their functions^3^.

Previously, chemically induced protein dimerization platforms such as rapamycin-based FKBP^4^, coumermycin-based GyrB^5^ and gibberellin-based^6^ systems have been employed to manipulate subcellular localizations of various proteins in both bacteria^7^ and eukaryotic^8^ cells. While relatively fast and robust, these methods lack the precise spatial targeting ability and are often irreversible on fast timescales^4^. In contrast, optogenetic-based systems use light-modulated binding of a photoreceptor either with itself or with its associated protein ligand and is reversible^9,10^; the application of light activation can be precisely defined in space to target specific cell subpopulations and even within a single cell’s volume^11–16^. Currently available light-induced protein-protein interaction systems have been extensively characterized in mammalian cell systems^3^ and have kinetic rates on timescales ranging from milliseconds to hours^17^. Despite the widespread use of optogenetic systems in eukaryotic cells, their use in bacteria is only beginning to emerge^3^. The major difficulty of their implementation resides in the bacteria cell’s 1000-fold smaller cellular volume compared to that of a mammalian cell^18^. Such a small cytoplasmic volume imposes strict requirements on the concentration and kinetic ranges of the optogenetic systems in order to achieve a high signal contrast and spatial resoltuion^18,19^.

One particular system, the CRY2/CIB1 system from *Arabidopsis thaliana*, can be induced by a low dose of blue light (peak activation wavelength at 468 nm) and exhibits rapid heterodimerization kinetics with tight^20^, but reversible^19^, binding. These properties make it an attractive candidate for use in model bacterial systems. The system is composed of the N-terminal domain (amino acids 1-170) of *At*CIB1 (CIBN) and the photolyase homology region^21^ (amino acids 1-498) of *At*CRY2 (CRY2)^19,22^. Here we show that this optogenetic system can be used to rapidly target proteins to major subcellular compartments in living *Escherichia coli* cells (**Fig. 1a**, left). We further demonstrate that such an optogenetically controlled system can be used to inhibit cytokinesis in living *E. coli* cells. Finally, we show that the association and dissociation kinetics of the system can be modulated by green light, adding a new dimension of control to suit different synthetic biological systems.

**Fig. 1.**
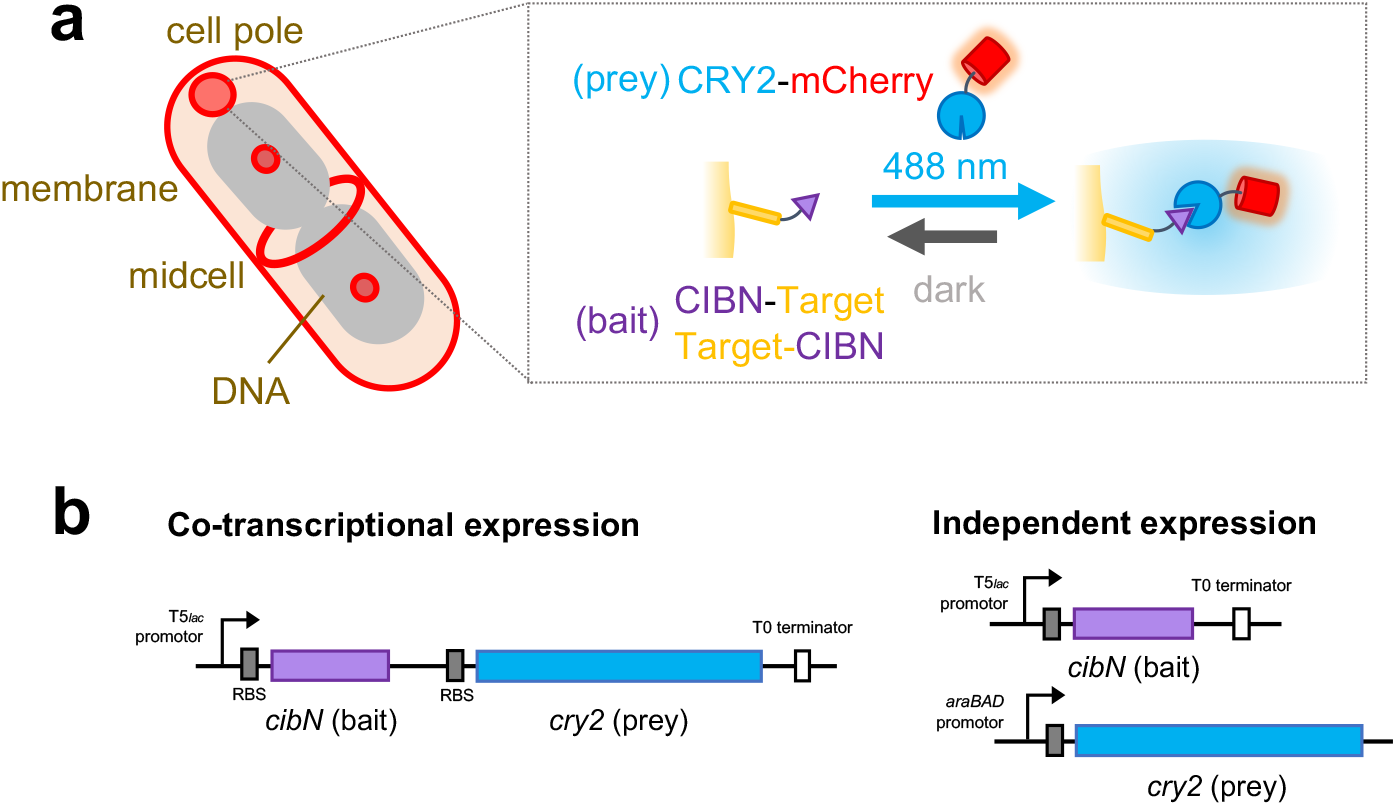
Light-induced recruitment scheme using CRY2 and CIBN. (**a**) CRY2 is targeted to different subcellular localizations in live *E. coli* cells: the chromosomal DNA, cell pole, inner membrane, and the midcell division plane. At each subcellular location, CIBN is fused to a bait protein at the target either at the C or N terminus, and CRY2 is fused to the N terminus of mCherry as a universal prey reporter for CIBN’s light (488 nm)-dependent association with CIBN-Target (inset box). The association is reversible in dark. (**b**) CIBN and CRY2 fusion proteins can be expressed co-transcriptionally (left) or independently (right) to suit different experimental needs.

## RESULTS

### General considerations for using the CRY2/CIBN system in *Escherichia coli* cells

Previous studies have shown that CIBN can be tagged at either its N- or C-terminus without impeding its binding to CRY2, whereas CRY2 functions the best with its N-terminus free^23^. Therefore, in the following experiments, we fused CIBN at its N- or C-terminus to an array of proteins that localize to a subcellular compartment of interest to serve as the bait for the recruitment platform and fused a fluorescent reporter, mCherry, to the C-terminus of CRY2 to serve as a universal prey (**Fig. 1a**, right). The use of the same CRY2-mCherry reporter to a variety of subcellular recruitment platforms allows us to compare recruitment kinetics between different systems.

Next, to attain the desired control of CRY2/CIBN expression levels inside small bacterial cells in a variety of experimental scenarios, we exogenously expressed the CRY2/CIBN fusion proteins either co-transcriptionally from a single plasmid where the two fusions were coupled under the control of a *lac-*inducible promotor, or independently from a two-plasmid system where the CRY2 and CIBN fusions were under the control of arabinose- and *lac-*inducible promoters, respectively (**Fig. 1b**). Whether the one- or two-plasmid expression system was employed depended on specific considerations for each experiment outlined below. We found that the flexibility in modulating the expression levels of the CRY2 and CIBN fusion proteins is important for minimizing light-independent background interaction^19^ while maintaining a fast light-dependent recruitment speed (discussed below).

### Rapid, reversible and light-dependent recruitment of cytoplasmic protein to chromosomal DNA

In bacterial cells, the nucleoid is the major subcellular location for all DNA-related processes. The ability to optically control chromosomal DNA topology (*e*.*g*., the formation of DNA loops or topological domains) or the binding of transcription factors on specific DNA sequences has proven useful to study transcription dynamics and gene expression^19,24,25^ in mammalian and bacterial cells^24,26–29^. Therefore, we chose the chromosomal DNA as the first platform to develop a light-dependent recruitment assay of cytoplasmic proteins.

To visualize the light-dependent chromosomal recruitment process, we used an *E. coli* strain harboring a 240X *tetO* array sequence inserted near *oriC* in its chromosome^30^ (**Fig. 2a**). The presence of tandem *tetO* arrays enhances the mCherry signal strength and facilitates the characterization of the recruitment kinetics. We then fused CIBN to the C-terminus of TetR, the tetracycline repressor that binds tightly to its cognate *tetO* operator sequence^31^ and co-transcriptionally expressed the TetR-CIBN and CRY2-mCherry reporter fusions exogenously from a single plasmid (**Fig. 1c**, left). The use of the coupled expression system helps to maintain a low and near equimolar expression of the two fusions proteins to best accommodate the 1:1 binding ratio of TetR-CIBN and CRY2-mCherry, effectively minimizing the background signal that would arise from excess CRY2-mCherry reporter. In the absence of the blue (488 nm) activation light, CRY2-mCherry remained homogenously distributed in the cytoplasm as expected for freely diffusing CRY2-mCherry in its unbound state (**Fig. 2b**, left). Upon exposure to 488 nm activation light pulses (30 ms pulses at 84.6 W/cm^2^ delivered every 5 s over a 200 s period), we observed rapid formation of CRY2-mCherry foci at locations representative of individual *oriC* sites in almost all cells in the view field (96 ± 1.3 %, μ ± s.e.m., mean ± standard error of the mean, *N* = 3 independent experiments of 1359 cells in total, **Fig. 2b**, right, **Fig. 2c**, top, **Supplementary Video 1**) across a large range of CRY2-mCherry expression levels in individual cells. Notably, the cytoplasmic distribution of CRY2-mCherry remained unchanged in control cells exposed to the 488 nm light but that did not contain the 240X *tetO* array (**Fig. 2c**, middle panel, **Supplementary Video 2**), or in cells that contained the *tetO* array but not exposed to the 488 nm activation light (**Fig. 2c**, bottom panel, **Supplementary Video 3**). Quantification of the CRY2-mCherry signal (percentage of fluorescence increase compared to that before activation) at foci positions over time showed that 90% recruitment was reached within 85 s (*τ*0.9 = 85 ± 9 s, μ ± s.e.m., *N* = 3 independent experiments of 442 cells in total, *τ*0.9 was used to show the extent of reaction completion as the curve was not well described by an exponential function, **Fig. 2d**). These results demonstrate that the formation of CRY2-mCherry foci is rapid and both light- and target-dependent.

**Fig. 2.**
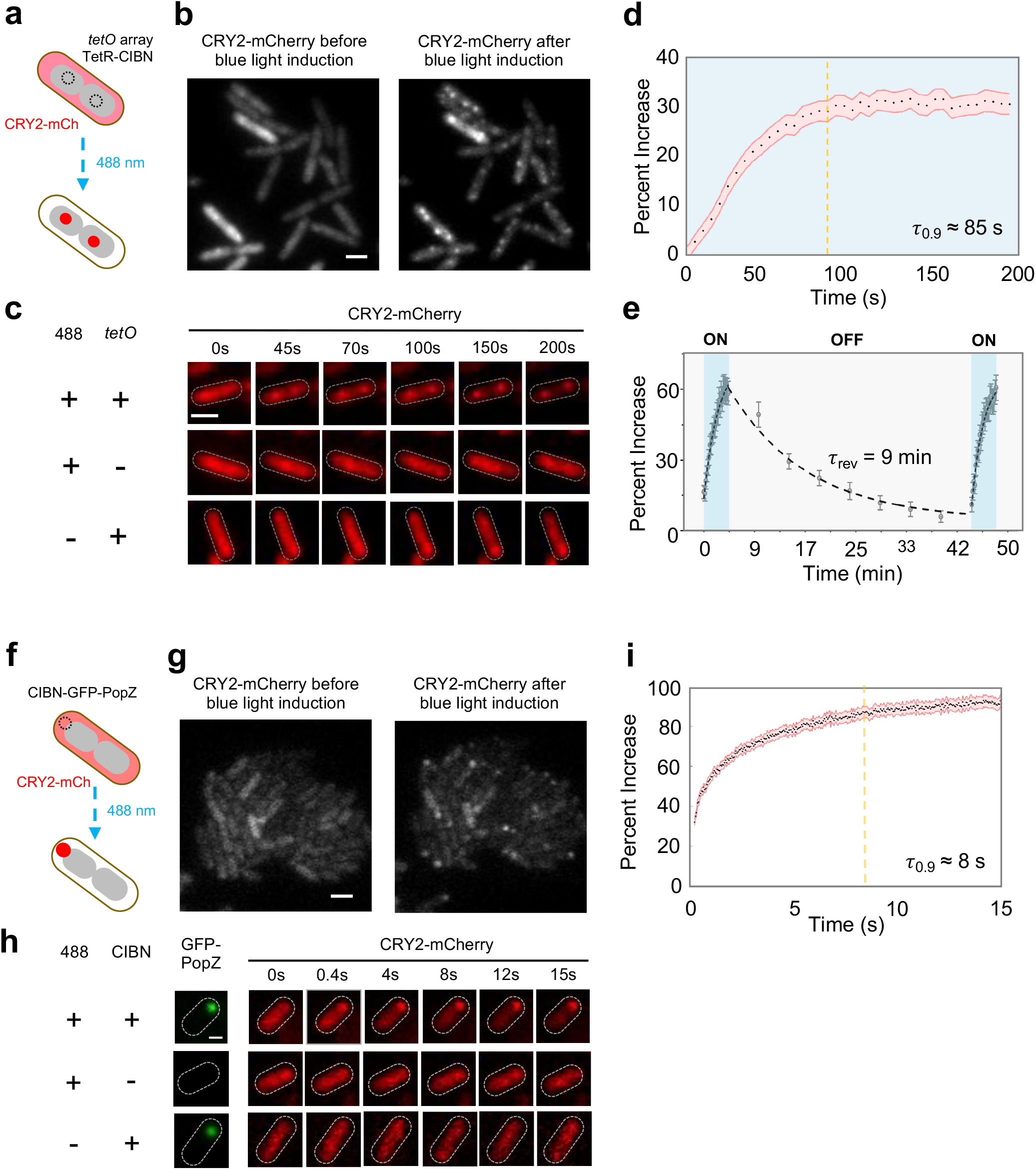
Rapid and reversible recruitment of cytoplasmic proteins to chromosomal DNA and Cell Pole. **(a)**, Schematic depicting the relocalization of cytoplasmic CRY2-mCherry to TetR-CIBN bound *tetO* sites near *OriC* to from foci after activation with blue light. (**b**), CRY2-mCherry is uniformly distributed throughout the cell prior to blue light exposure *(left panel*). After blue light exposure CRY2-mCherry rapidly relocalizes to form foci (*right panel*). (**c**), Single cell time course images demonstrating that the recruitment of CRY2-mCherry to DNA foci occurs only when blue light and the *tetO* array are both present (*top row*) but not in the absence of the *tetO* target (*middle row*) or blue light activation (*bottom row*). (**d**), Averaged percent increase of CRY2-mCherry signal at DNA foci (*N* = 1359 cells) demonstrating that 90% recruitment is reached within 85 seconds. (**e**), Averaged percentage increase of CRY2-mCherry signal at DNA foci after activation with blue light (*first blue section*), relaxation in the dark (*middle grey section*), and after a second activation sequence (*second blue region*) demonstrating that CRY2/CIBN disassociation at DNA foci is reversible with a time constant of ∼9 minutes. (**f**), Schematic depicting the relocalization of cytoplasmic CRY2-mCherry to CIBN-GFP-PopZ foci at the cell pole after blue light activation. (**g**) CRY2-mCherry is uniformly distributed throughout the cell prior to blue light exposure *(left panel*). After blue light exposure, CRY2-mCherry rapidly relocalizes to the cell poles and form foci (*right panel*). (**h**) Single cell time course images demonstrating that the recruitment of CRY2-mCherry to the cell pole occurs only when both blue light and the CIBN are present (*top row*) but not in the absence of the CIBN target (*middle row*) or blue light (*bottom row*). (**i**), Averaged percent increase of CRY2-mCherry signal at cell pole foci (*N* = 315 cells) demonstrating that 90% recruitment is reached within ∼8 seconds.

In addition to its rapid association kinetics, the CRY2/CIBN system is also reversible in the absence of blue light^19^. When we allowed the CRY2/CIBN system to relax in the absence of blue light following the complete localization of CRY2-mCherry to *oriC* foci (**Fig. 2e**, first blue shaded region), we observed an almost complete reversion of CRY2-mCherry foci back to a uniform cytoplasmic distribution after ∼ 40 minutes (**Fig. 2e, Supplementary Figure 1**). The decay curve was well described by an exponential function with a relaxation time constant of ∼ 9 minutes (r_rev_ = 9 ± 5 min, µ ± s.e.m., *N* = 3 independent experiments of 150 cells, **Fig. 2e**, shaded grey region). This relaxation time scale is consistent with the reversion times measured previously in mammalian cells^17,19^. Additionally, relaxed cells could be activated once again^17,19^ to re-form the same *oriC* foci with the same recruitment rate and final plateau (**Fig. 2e**, second blue shaded region and **Supplementary Video 4**). In summary, these results clearly demonstrate that the CRY2/CIBN system can be used to induce reversible recruitment of a cytoplasmic protein to chromosomal DNAs rapidly and efficiently in live *E. coli* cells.

### Rapid, light-dependent recruitment of cytoplasmic protein to cell pole

In bacteria, asymmetric cell division is often achieved by partitioning proteins to each incipient daughter cell *via* cell pole localized protein scaffolds^32^. In symmetrically dividing bacteria such as *E. coli*, the two poles are also differentiated in age, and proteins can be targeted specifically to the old cell pole to produce a dimorphic cell population^33,34^. Furthermore, the cell pole can be engineered into an optically controlled apical protein sink to present an inert space away from the cytoplasmic environment^35^.

To develop a light-triggered cell pole recruitment platform, we took advantage of a protein called PopZ from *Caulobacter crescentus*. PopZ forms a stable, liquid droplet-like matrix enriched at the old cell pole when heterologously expressed in *E. coli* cells^36– 38^ (**Fig. 2f**). To visualize the cell pole localization of PopZ, we fused CIBN-GFP to the N terminus of PopZ and placed the fusion gene under a *lac*-inducible promoter on a plasmid (**Fig. 1b**, right). After optimizing the expression of the PopZ fusion, we observed large, stable CIBN-GFP-PopZ foci at the cell poles of nearly every cell (**Supplementary Fig. 2**). We then expressed the CRY2-mCherry fusion under the control of an arabinose-inducible promoter from a separate vector (**Fig. 1b**, right). Here we used the two-plasmid system for independent expression control of the two fusion proteins, because the liquid droplet nature of the PopZ foci can accommodate a very high expression level of CIBN-GFP-PopZ, which ensures a stable cell pole localization and to provide a sufficient number of binding sites for CRY2-mCherry.

As shown in **Fig. 2g** and **Supplementary Video 5**, following a single 100 ms blue light activation pulse (*P*_488_ *=* 8.5 W/cm^2^), we observed rapid formation of CRY2-mCherry foci at the cell poles. Importantly, in cells exposed to blue light only expressing GFP-PopZ, but not CIBN-GFP-PopZ (**Fig. 2h**, middle panel and **Supplementary Video 6**), or in cells without blue light activation (**Fig. 2h**, bottom panel and **Supplementary Video 7**), we observed unchanged, homogenous distribution of CRY2-mCherry signal in the cytoplasm throughout the time course of the experiment. Quantification of the percent increase in CRY2-mCherry signal at the cell poles revealed that 90% recruitment was achieved within ∼8 s (*τ*0.9 = 8.3 ± 0.9 s, μ ± s.e.m., *N =* 3 repeats of 315 cells in total, **Fig. 2i**). Note that in this system, the recruitment of cytoplasmic CRY2-mCherry was sped up to ∼10-fold compared to that in the DNA recruitment assay. The faster kinetics are most likely driven by the higher expression levels of both CIBN-GFP-PopZ and CRY2-mCherry in the system.

### Recruiting protein cargo to the inner membrane using CRY2/CIBN

In many cell types, the inner membrane (IM) is the site where a diverse array of signaling pathways begin. In bacteria, the IM also serves as the site of assembly for a number of macromolecular structures including the divisome^39^, the flagellum^40^, and chemotaxis receptor clusters^41^. In mammalian cells, synthetic activation of membrane-based signaling cascades be achieved using cytoplasmic CRY2-tagged proteins that are recruited to the plasma membrane by membrane-anchored CIBN^16,19,23,42,43^. To develop a light-dependent membrane recruitment assay in bacteria, we fused CIBN-GFP to an amphipathic helix from *Bacillus subtilis* (*Bs*MTS)^44^ and expressed it from a *lac-*inducible promotor independently from the arabinose-inducible CRY2-mCherry on a second plasmid. Here we used an exogenous membrane anchor to avoid potential interference with endogenous *E. coli* membrane proteins. *Bs*MTS-GFP-CIBN showed uniform membrane localization as expected (**Supplementary Fig. 3a**, left). Upon blue light activation (50 ms pulses at 84.6 W/cm^2^ delivered every 5 s), we observed the formation of dynamic CRY2-mCherry puncta that co-localized with CIBN-GFP-*Bs*MTS puncta along the IM (**Supplementary Fig. 3b**, & **Supplementary Video 8**). The distributions of these puncta were distinct from the near-uniform distribution of CIBN-GFP-*Bs*MTS along the IM surface prior to activation with blue light (**Supplementary Fig. 3a**, left), or that of directly membrane-targeted CRY2-GFP-BsMTS (**Supplementary Fig. 3a**, right). We reason that the puncta formation along the IM of *Bs*MTS-CIBN-GFP and CRY2-mCherry upon light activation is likely indicative of the formation of individual membrane-tethered CIBN-CRY2 hetero clusters as previously predicated^12,23,45^. Interestingly, despite a large number of strategies employed (**Supplementary Table 1**, reference ^46^), we did not observe uniform co-localization of CRY2 and CIBN on the IM of *E. coli* cells in contrast to what was observed previously in mammalian cells^12,19^. Further experiments are required to investigate this phenomenon.

### Rapid, light-dependent recruitment of cytoplasmic proteins to the midcell

In almost all bacteria studied to date, the essential process of cytokinesis is mediated by the formation of a large macromolecular complex, termed the divisome, at the future division plane of the cell^47^. In *E. coli*, the divisome is comprised of more than 30 different proteins, which are recruited to the midcell by the essential tubulin homologue, FtsZ^39,48^. FtsZ polymerizes at the midcell to assemble into a ring-like structure, and new data suggests that the dynamics of the Z-ring are essential to coordinate the action of the divisome for cell wall constriction and septum morphogenesis during cell division^49–51^. Achieving a means to rapidly and reversibly deliver proteins to the divisome and/or modulate the assembly of the divisome at different cell division stages would provide a new way to probe the dependence of different divisome proteins and their functions on the assembly state of the divisome specifically at the division site^52^.

To achieve optically controlled delivery of proteins to the cell division plane, we first tested the ability to recruit CRY2-mCherry to the Z-ring by creating a C-terminal fusion of CIBN to ZapA (**Fig. 3a**). ZapA is a conserved Z-ring-associated protein, which has served as a faithful Z-ring marker in previous studies^53–56^. Similar to the pole-recruitment platform, we expressed the ZapA-CIBN fusion independently from the CRY2-mCherry reporter using the two-plasmid expression system so that their expression levels could be optimized separately. Upon exposing cells to a single 100 ms pulse of 488 nm activation light (*P*_488_ *=* 8.5 W/cm^2^), we observed rapid midcell localization of CRY2-mCherry fluorescence (*τ*_0.9_ = 9.3 ± 0.15 s, μ ± s.e.m., *N* = 3 repeats of 443 cells in total, **Fig. 3b-d, Supplementary Video 9**). Similar to what was observed in the chromosomal DNA and cell pole recruitment assays, the midcell localization of CRY2-mCherry was dependent on the blue activation light (**Fig. 3c**). These results demonstrate that cytoplasmic proteins can be recruited to the Z-ring at the midcell rapidly and specifically using the CRY2/CIBN system.

**Fig. 3.**
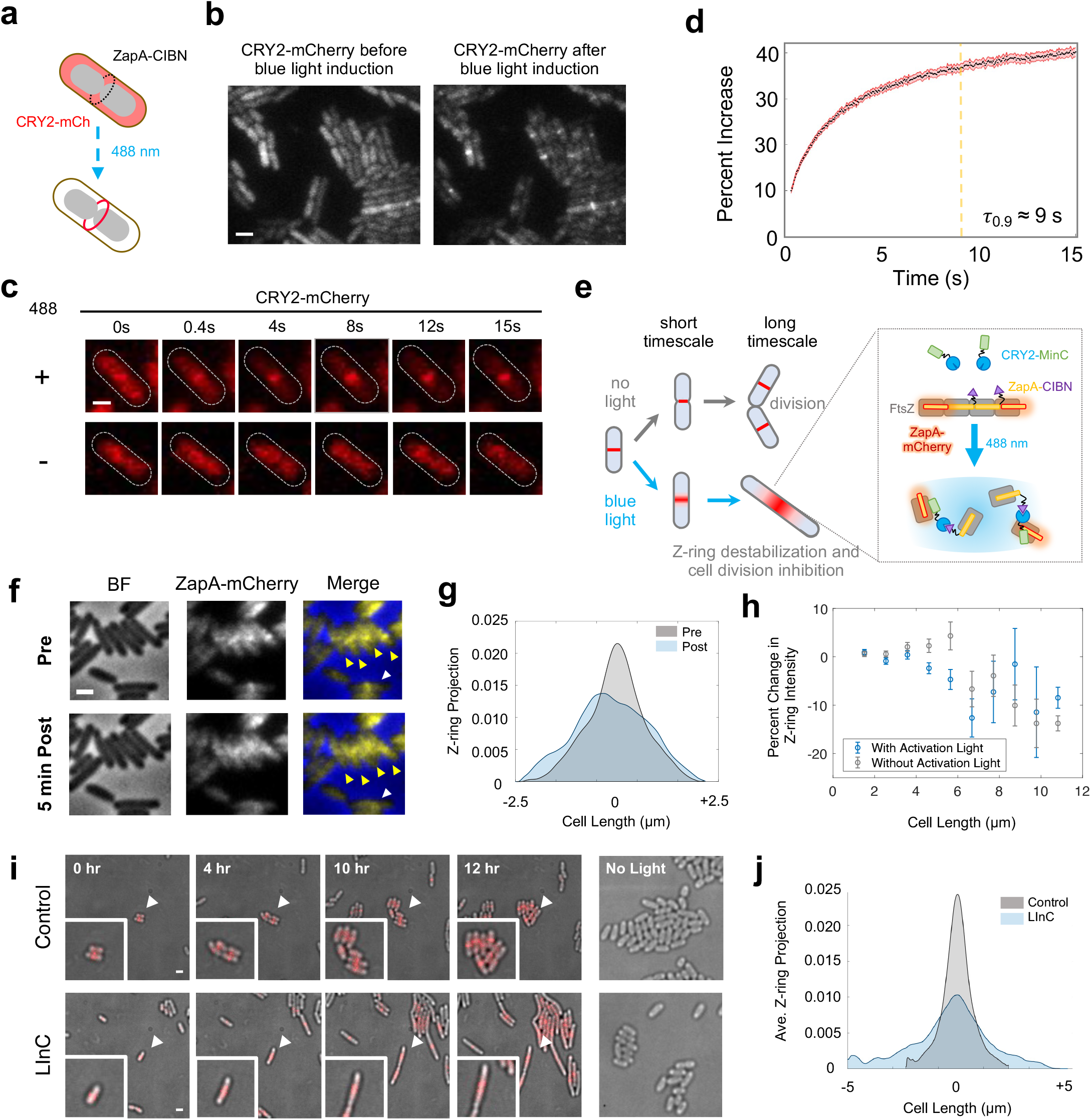
Recruitment of cytoplasmic protein to cell division plane and a Light-induced Inhibition of Cytokinesis (LInC) Assay. (**a**), Schematic depicting the relocalization of cytoplasmic CRY2-mCherry to the ZapA-CIBN ring present at midcell after induction of CRY2/CIBN binding with blue light. (**b**), An example image of cells showing CRY2-mCherry’s diffusive cytoplasmic localization before blue light activation (*left*) and midcell localization after blue light activation (*right*). (**c**), Single cell time course images demonstrating that CRY2-mCherry recruitment to midcell occurs only after activation with blue light. (**d**), Averaged percent increase of CRY2-mCherry signal at midcell demonstrating that 90% recruitment is reached within 9 seconds (*N* = 443 cells). (**e**), Schematic depicting the LInC assay on both short and long time scales. Recruitment of CRY2-MinC to Z-rings *via* ZapA-CIBN by blue light activation results in instant destabilization of the Z-ring at a short time scale and cell division inhibition at a long time scale. In the absence of blue light cells grow and divide like WT cells. (**f**), **Short timescale LInC Assay**. Cells harboring the LInC system exhibited diffusive Z-rings (arrow heads) after blue light activation delivered every 10 s for 5 min. The projected fluorescence intensity of ZapA-mCherry (Z-ring) along the cell long-axis before (grey) and after a 5-minute blue light induction (blue) for the cell labeled with a white arrow in **f** is plotted in (**g)** to demonstrate the decondensation of the Z-ring. (**h**), Short timescale Z-ring destabilization by the LInC system is dependent on cell length. The percent reduction of ZapA-mCherry intensity (μ ± s.e.m., **Supplementary Table 2**) at midcell in cells expressing the LInC system before and after a 5-minute period of blue light activation (blue datapoints) or darkness (grey datapoints) plotted as a function of cell length. (**i**), **Long timescale Linc assay**. Time-lapse imaging shows that cells ectopically expressing CRY2-MinC only (Control, *top row*) grew and divided normally while cells expressing both CRY2-MinC and ZapA-CIBN (LInC, *bottom row*) filamented when both were exposed to 100 ms pulses of blue and green light every 5 minutes for a 12-hour period. Cells harboring only CRY2-MinC (right panel, *top*) or both CRY2-MinC and ZapA-CIBN (right panel, *bottom row*) but were not exposed to blue light divided normally during the same period (right most panel). White arrows indicate the position of cells that are enlarged in the insets. (**j**), Smoothed and averaged long axis projections of the Z-ring intensity measured by ZapA-mCherry at the 12-hour timepoint after blue light activation demonstrated significant widening of Z-ring in cells harboring the LInC system (FWHM = 221 nm, *N* = 16 cells) compared with those in the Control cells (FWHM = 102 nm, *N* = 33 cells).

### Light-dependent destabilization of the Z-ring and inhibition of cell division

We next used the midcell recruitment platform to develop a Light-induced Inhibition of Cytokinesis (LInC) assay in live *E. coli* cells (**Fig. 3e**). In the LInC assay, we kept the same ZapA-CIBN fusion as the bait, but fused a Z-ring antagonist, MinC, to CRY2 to generate a CRY2-MinC fusion. MinC inhibits FtsZ polymerization *in vitro* and prevents Z-ring assembly at the cell poles by forming a high to low concentration gradient from the cell pole to the midcell with its membrane binding partner ATPase MinD^57^. When highly overexpressed by itself, MinC antagonizes Z-ring assembly at the midcell as well^57–59^. We verified that the CRY2-MinC fusion is fully functional as it complemented a *minC* deletion background (**Supplementary Fig. 4**). Therefore, by recruiting CRY2-MinC to FtsZ-associated ZapA-CIBN, we expect to increase the local concentration of MinC at the Z-ring, which may result in the destabilization of the Z-ring and subsequent inhibition of cell division (**Fig. 3e**). To visualize the progress of Z-ring destabilization in the LInC assay, we tagged the endogenous chromosomal copy of ZapA with mCherry as the Z-ring marker^53,54^.

We first established a growth and expression conditions under which cells ectopically expressing the LInC system (ZapA-CIBN and CRY2-MinC) exhibited a near-WT midcell Z-ring morphology and underwent successful cell division cycles in the absence of blue light (**Supplementary Note 1**). Notably, these cells were moderately longer than uninduced cells due to the presence of higher than WT CRY2-MinC concentrations (*L =* 3.04 ± 0.17 μm, μ ± s.e.m., *N* = 3 repeats of 638 cells in total compared to *L* = 2.45 ± 0.024 μm, μ ± s.e.m., *N* = 3 repeats of 223 cells in total for uninduced cells). We then imaged the Z-ring morphology of these cells before and after exposing them to 100 ms blue light pulses (*P*_488_ *=* 84.6 W/cm^2^) every 10 seconds for a period of 5 minutes (**Fig. 3f**). Before the activation light exposure, we quantified that on average ∼ 46% of all the cells harbored clear ZapA-mCherry localization at midcell, consistent with previous reports that the Z-ring only assembles after a significant portion of the chromosome is segregated^60^. About 52% of these cells (51.9 ± 0.4%, μ ± s.e.m., *N =* 1658 cells) exhibited significantly broader and more diffuse ZapA-mCherry midcell fluorescence upon exposure to blue light compared to that prior to light exposure (**Fig. 3f** & **g**), indicating the destabilization of the Z-ring. In control cells with only the CRY2-MinC fusion or without blue light activation, we observed significantly lower, ∼ 37% (36.5 ± 0.8%, μ ± s.e.m., *N =* 379 cells) or ∼ 44%, (43.8 ± 0.4%, μ ± s.e.m., *N =* 1465 cells), cells exhibiting diminished ZapA-mCherry midcell fluorescence after a 5 min period, indicative of the natural disassembly process of the Z-ring during cell division. Interestingly, the extend of the Z-ring destabilization appeared to be dependent on the cell division stage (**Fig. 3h, Supplementary Table 2**). We observed that after Z-ring assembly, cells of relatively short lengths (indicative of earlier cell division stages) showed significant destabilization of the Z-ring (**Fig. 3h**, blue) compared to cells of longer lengths, or cell of similar lengths but without the activation light (**Fig. 3h**, grey). Previous studies have shown that when the divisome matures and cell wall constriction initiates, the Z-ring becomes denser and denser until it finally and rapidly disassembles toward the end of the cell cycle^61,62^. Therefore, it is possible that the LInC assay destabilizes the Z-ring effectively only before the Z-ring becomes too dense to be effectively depolymerized by MinC.

While we were able to achieve efficient and rapid Z-ring destabilization in cells judged by their widened Z-ring morphology and the reduction in the midcell localization percentage of FtsZ, it remained unclear whether these destabilized Z-rings were indeed defective in cell division. To address this, we monitored cell division for a long period of time in cells harboring the LInC system after induction of the system with blue light. As shown in **Fig. 3i**, when we subjected LInC cells to 100 ms pulses of 488-nm activation light (*P*_488_ *=* 8.5 W/cm^2^) and 561-nm imaging light (*P*_561_ *=* 8.7 W/cm^2^) every 5 minutes for a twelve-hour period, we observed that the majority of these cells ceased division and grew into long filaments instead. On average, ∼ 76% (*N* = 33 cells) were unable to divide during the 12-h period. Additionally, the average division time (*τ*_LInC_ = 355 ± 120 min, μ ± s.e.m., *N* = 3 repeats of 33 cells in total) of the remaining ∼ 24% cells that were able to divide eventually was nearly 1.5 times longer than the mean division time of the control group where the ZapA-CIBN bait fusion was absent (*τ*_control_ = 232 ± 54 min, μ ± s.e.m., *N* = 3 repeats of 31 cells in total). These observations indicated significant deficiency of LInC cells in division. The inhibited cell division also correlated with significantly destabilized Z-rings—in all long, filamentous cells, the midcell fluorescence of ZapA-mCherry exhibited a wider spread, in contrast to the sharp midcell localization prior to light exposure (**Fig. 3j**). Note that the inhibition of cell division in these cells was not caused by photodamage due to the continuous illumination of the 488-nm activation light and the 561-nm imaging light, because cells under the same induction and illumination condition but ectopically expressing only CRY2-MinC grew and divided as wild type cells (**Fig. 3i**, top row rightmost panel). Furthermore, the inhibition was 488-nm light-dependent because cells expressing both CRY2-MinC and ZapA-CIBN in the absence of the blue light grew normally as WT cells (**Fig. 3i**, bottom row rightmost panel). Taken together, these results show that the CRY2/CIBN system can be used to destabilize the Z-ring rapidly and inhibit cell division in living *E. coli* cells in a light-dependent manner.

### CRY2/CIBN complex dissociation and activated CRY2 concentration are sensitive to green light

Previous studies have shown that the light sensitivity of CRY2 is imparted by its flavin adenine dinucleotide (FAD) cofactor through a photocycle^52,53^ (**Fig. 4a**). In this cycle, the redox state of CRY2’s FAD is thought to mediate a conformational change in CRY2 to form a signaling-active state that is competent to bind CIB^63^. Briefly, the fully oxidated FAD^ox^ cofactor is in the resting inactive state. Following photoexcitation by blue light, FAD^ox^ accepts an electron from a nearby tryptophan followed by protonation from an aspartic acid to become the semi-reduced, stable semiquinone radical (FADH°). This redox state is thought to trigger a conformational change in CRY2 (denoted as CRY2*) that allows for its binding to CIB^64,65^. To complete the photocycle, semi-reduced FADH° is reduced slowly to become FADH^-^, reversing the conformation change, after which it is oxidated rapidly back to its fully-oxidated resting state (FAD^ox^). Previous studies have indicated that the semiquinone form FADH° is the only green-absorbing species in FAD’s photocycle and that green light can deplete FADH° *in vitro*^65^. In short-day entrained plants, green light leads to a delay in flowering, likely due to the increased reduction of FADH° to FADH^-^ under green light exposure^54,55^. However, previous characterization of CRY2/CIBN complex formation *in vivo* in heterologous expression systems showed no sensitivity to green light^19^.

**Fig. 4.**
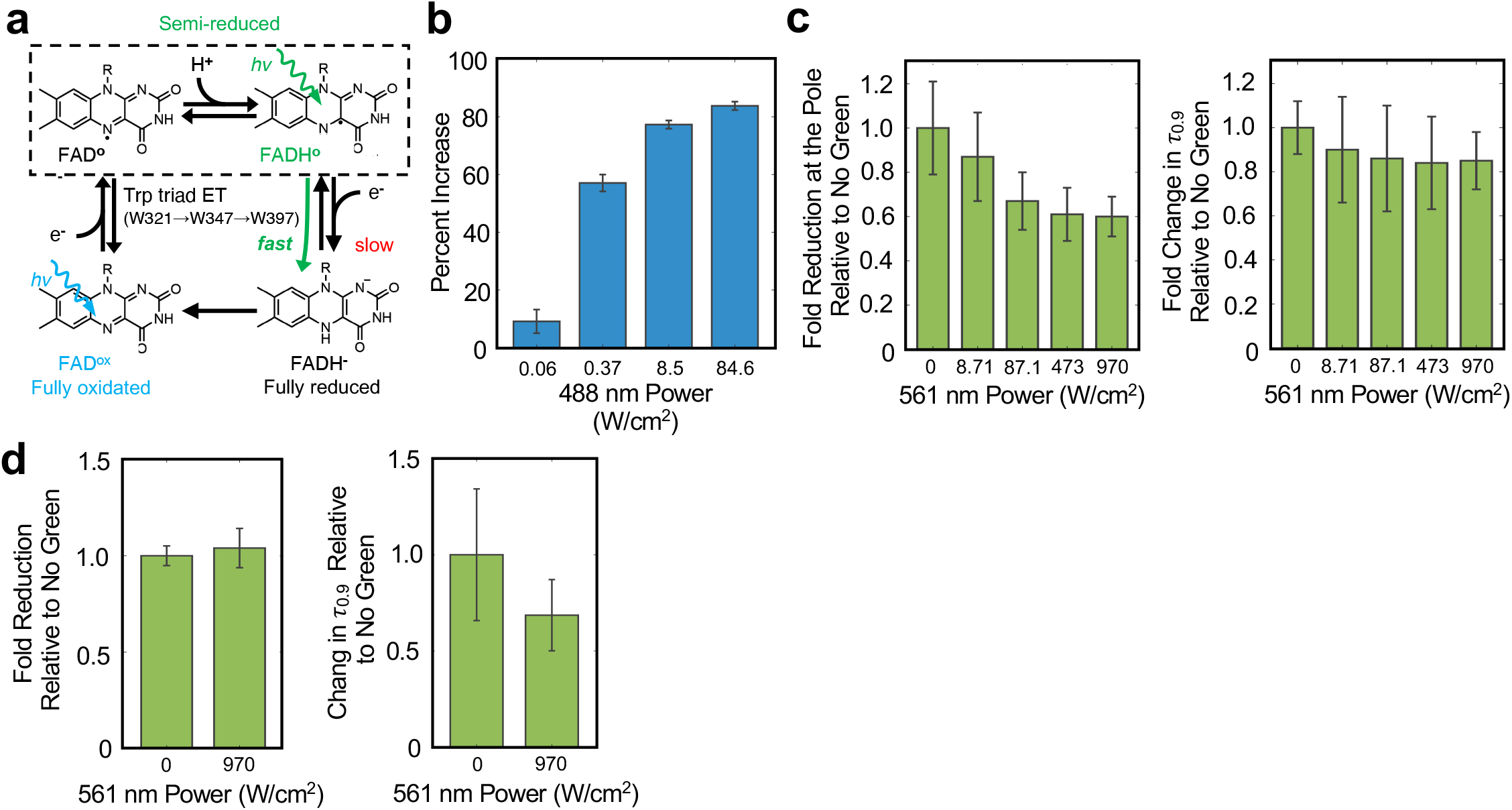
Green (561 nm) modulates the association and dissociation of CRY2-CIBN complex. (**a**), Schematic of CRY2 photoactivation pathway highlighting the redox states of its flavin (FAD) cofactor along the path. Breifly, the fully oxidized FAD^ox^ is photoexcited by a blue light photon and accepts an electron from the nearby tryptophan (Trp397) to form a stable, semi-reduced neutral FADH°. This formation induces a conformational change in CRY2, which allows it to bind to CIBN. In the absence of green light, the semi-reduced FADH° slowly decays to a fully reduced state (FADH^-^). As FADH° is the only species in the cycle that can absorb green light, it is likely that green light speeds up the further reduction of FADH° to FADH^-^,which prevents CRY2 to associate or remain associated with CIBN. (**b**), The percent enrichment of cell pole-recruited CRY2-Halo shows a stepwise dependence on the intensity of the 100ms blue (488 nm) light pulse used to trigger CRY2/CIBN complex formation. (error bars = s.d.) (**c**), The maximal fold enrichment of cell pole-recruited CRY2-Halo under varying green light intensities relative to that in the absence of green light shows a green light dependent reduction in a green-light dependent manner (*left*, error bars = s.d.), while the 90% recruitment time (τ_0.9_) under varying green light intensities relative to that in the absence of green light was not significantly altered (*right*, error bars = s.d.) indicating that green light can modulate the levels of activated CRY2 available for complex formation with CIBN. (**d**), The change in the final fold reduction of cell pole-recruited CRY2-Halo dissociation under green light exposure relative to that in the absence of green light showed little change (*left*, error bars = s.d.), while the 90% recruitment time (τ_0.9_) under green light exposure relative to that in the absence of green light showed a green light dependent reduction (*right*, error bars = s.d.) indicating that green light can modulate the dissociation rate of the CRY2/CIBN complex.

We reason that if the association/dissociation kinetics of the CRY2/CIBN are indeed sensitive to green light, we could use green light (561 nm) in combination with blue activation light to fine tune the kinetic behavior of the CRY2/CIBN system. To investigate this possibility, we swapped the mCherry reporter with Halo^66^, which can be covalently labeled with a bright, photostable far-red organic dye^67^ (JF646), allowing us to avoid using green light to image the reporter. We verified that the extent of CRY2/CIBN association indicated by the percentage increase in cells expressing the Halo-labeled system exhibited a blue-light activation dose dependence (**Fig. 4b**) as expected^19^, and that the final association plateau could be further increased using subsequent multiple pulses of activation to recruit additional CRY2-Halo to the pole (**Supplementary Fig. 5**). These results indicate that CRY2-Halo can be activated in a similar manner as CRY2-mCherry by a 100 ms blue pulse, and that the final pole-enrichment of CRY2 is dependent on the available level of activated CRY2, which can be modulated by the intensity of the activation pulse.

Next, to investigate the influence of green light on CRY2/CIBN association, we monitored CRY2/CIBN complex formation at the cell poles following a single 100 ms pulse of 8.5 W/cm^2^ blue activation light followed by a continuous pulse of green light of different intensities (561 nm, from 8.71 to 970 W/cm^2^) accompanying the imaging 647 nm light (50 ms exposure for 15 seconds). We found that while the time constant *τ*_0.9_ values did not appear to change significantly, the amount of CRY2 recruited to the cell pole decreased significantly with increasing green light intensity (**Fig. 4c**, *right and left panels* respectively, **Supplementary Fig. 6, Supplementary Table 3**). At the highest green light intensity (970 W/cm^2^), the pole-recruited CRY2 fraction was reduced to ∼ 50% compared to that in the absence of the green light. These observations indicate that green light can potentially drive activated CRY2* back to the inactivated state even before it forms a complex with CIBN, hence modulating the level of activated CRY2* and subsequently the amount of CRY2*-CIBN complex formation in live *E. coli* cells.

To investigate whether green light also modulates CRY2/CIBN dissociation kinetics, we light-induced CRY2-CIBN complex formation using blue light and then monitored the dissociation kinetics of the complex in the presence or absence of green light (100 ms pulses at 561 nm light 473 W/cm^2^ delivered every 3 min). We found that the time constant of the dissociation reaction decreased ∼ 30%, from 18.3 ± 8.18 min (*N* = 4 repeats with 91 cells in total) to 12.7 ± 5.70 min (μ ± s.e.m., *N* = 4 repeats with 62 cells in total) in the presence of the green light, while the final plateau did not change dramatically (**Fig. 4d**, *left and right panel* respectively, **Supplementary Fig. 7, Supplementary Table 3**). This observation suggests that the green light can also act upon activated CRY2* in complex with CIBN to drive it back to the inactivated from, effectively speeding up the dissociation of the CRY2*-CIBN complex, likely through the photoreduction of FADH° within the CRY2*-CIBN complex.

## Discussion (<200 words)

The landscape of optogenetic systems to induce protein-protein interactions inside living cells is constantly growing^3,68^. Optogenetic systems are especially useful for live-cell imaging studies due to their rapid induction, reversible binding and spatial precision^69,70^. Here we demonstrate that the *Arabidopsis thaliana* CRY2/CIBN system (**Fig. 1a**), used extensively in mammalian cell systems, can be applied in living *E. coli* cells to target a protein cargo to different subcellular structures including the chromosomal DNA, cell pole, inner membrane and the division plane at the midcell (**Fig. 1b**). We found that the expression levels of CRY2/CIBN system in bacteria cells can be varied to achieve recruitment halftimes ranging from the seconds to minutes timescales similar to those observed in mammalian cells^19,23^. We then demonstrate that the CRY2/CIBN system can also be implemented to rapidly destabilize the cytokinetic ring and inhibit cell division in *E. coli* (**Fig. 3**), providing a rapid and spatially specific tool to manipulation bacterial cell division.

During our characterization of the CRY2/CIBN system, we found that both the association and dissociation of CRY2 with CIBN can be modulated by green light (**Fig. 4c**). This effect not only enables another layer of system tuning akin to the blue light dose-dependent activation behavior (**Fig. 4b**) that we and others^19,25^ have observed, but also provides direct evidence that green light inactivates CRY2 likely by acting on the semi-reduced FADH° cofactor directly to reverse the active conformation of CRY2, hence lowering the amount of CRY2* available for complex formation and also speeding up the dissociation kinetics of previously formed CRY2*-CIBN complex. These results are consistent with previous experiments where green light exposure was found to rapidly convert CRY2’s cofactor from FADH° to FAD_ox_ *in vitro* and inhibit CRY2/CIB1 activity *in vivo*^65^.

In mammalian cells the CRY2/CIBN system has already been employed to optogenetically control cellular pathways including Cre recombination^71^, Raf/MEK/Erk^23^, TGF-β^72^, NF-*κ*B^15^, Wnt/β-catenin^11^, and RhoA^11^. Most of these systems involve the binding of two proteins to initiate a signaling cascade. In bacteria, we envision that the CRY2/CIBN system can be similarly applied to a large number of processes to enable optogenetic control. For example, most of the two-component signaling (TCS) systems^73^ in bacteria employ a histidine kinase that, upon stimulation, will recruit and phosphorylate its cognate response regulator resulting in the activation of downstream signaling^73^. These protein pairs could be tagged and controlled in a similar manner as done in eukaryotes to allow for the pathway regulation in real-time. Furthermore, a number of pre-existing chemically inducible systems could be transformed into optogenetic systems by modifying their dimerization platforms. Potential candidate systems could include split ClpXP adaptors for optical control of protein degradation^7^, split T7 RNAP for optical control of gene expression^24^ and split Cas9 for optical control of gene editing^71^.

### METHODS

### Bacterial strains, plasmids and growth conditions

All *E. coli* strains used for the cell pole recruitment, Z-ring recruitment, and cytokinesis inhibition experiments were derivatives of BW25113 (CGSC #7636) strain and were grown in M9+ Glucose minimal media (recipe is provided in **Supplementary Table 4**). The DNA recruitment background strain was previously constructed by the Sherratt Lab^30^ (CGSC #12294) and was grown in EZ Rich Defined Media (EZRDM, Teknova Bio.) as described in **Supplementary Table 4**. Antibiotic requirements are specified in **Supplementary Table 4** and the corresponding concentrations for each were chloramphenicol (150μg/mL), streptomycin (50μg/mL) and gentamycin (5μg/mL). Inducer concentrations for respective strains and plasmid combinations are outlined in **Supplementary Table 4** and described in more detail below. All bacterial cultures were grown in tubes covered in foil to minimize contact with ambient light. Plasmids expressing exogenous CRY2/CIBN fusion proteins with their associated sequences are described in **Supplementary Table 4**.

### Recruitment of CRY2-mCherry to the DNA *via* TetR-CIBN

Cultures were started from a single colony of the strain RM187 and grown in 6mL EZ Rich Defined Medium (EZRDM, Teknova Bio.) containing chloramphenicol and gentamycin at 25ºC with shaking until the cells entered early log-phase (OD600 between 0.1 and 0.2, approx. 19 hours). When the culture entered early log-phase the cells were induced with 60μM IPTG and supplemented with 240nM anhydrous-tetracycline (ATC) to avoid a replication block induced by binding of TetR-CIBN similarly as previously described. The cells were induced for 2 hours at 25ºC with shaking. At the end of the induction period the cells were centrifuged for 10 minutes at 4110k rpm and the pellet was resuspended in 6mL of EZRDM containing chloramphenicol and 240nM ATC and grown for an additional 30 minutes at 25ºC while shaking after which it was, again, centrifuged for 10 minutes at 4110k rpm and resuspended in 6mL EZRDM containing chloramphenicol and gentamycin and allowed to grow an additional 2.5 hours at 25ºC while shaking to allow the mCherry fluorophore to mature. Following the outgrowth period, the cells were prepared for imaging by centrifuging 1mL of the culture at 13k rpm for 1 minute and then resuspending the pellet in 50uL of fresh EZRDM. Following this, 0.5uL of the resuspension was then placed on a gel pad and the pad and prepared for imaging as specified in the “Live cell imaging” section below.

### Recruitment of CRY2-mCherry to the cell pole *via* CIBN-GFP-PopZ and the Z-ring *via* ZapA-CIBN

Cultures were started from a single colony of the strain of interest and grown in M9+ Glucose minimal medium containing chloramphenicol and streptomycin at 37ºC overnight until the cells reached stationary phase. The overnight culture was then diluted 1:100 in fresh of M9+ Glucose containing chloramphenicol and streptomycin and was allowed to grow at 37ºC until it reached log-phase (OD600 between 0.1 and 0.2, approx. 3 hours) at which point the culture was induced with 0.4% arabinose and either 40μM IPTG (ZapA fusion) or 100μM IPTG (PopZ fusion). Following a 1-hour induction at 37ºC the cells were prepared for imaging by centrifuging 1mL of the culture at 13k rpm for 1 minute and then resuspending the pellet in 70uL of fresh M9+ Glucose. Following this, 0.5uL of the resuspension was then placed on a gel pad and the pad and prepared for imaging as specified in the “Live cell imaging” section below. The cells on the gel pad were then assembled into the imaging chamber and allowed to equilibrate on the microscope in the dark for 2 hours at ambient room temperature (RT).

### Recruitment of CRY2-HaloTag to the cell pole *via* CIBN-GFP-PopZ using variable green (561nm) light

Cultures were started from a single colony of the strain of interest and grown in M9+ Glucose minimal medium containing chloramphenicol and streptomycin at 37ºC overnight until the cells reached stationary phase. The overnight culture was then diluted 1:100 in fresh of M9+ Glucose containing chloramphenicol and streptomycin and was allowed to grow at 37ºC until it reached log-phase (OD600 between 0.1 and 0.2, approx. 3 hours) at which point the culture was induced. Following a 1-hour induction at 37ºC the cells were prepared for imaging by centrifuging 1mL of the culture at 13k rpm for 1 minute and then resuspending the pellet in 100uL of fresh M9+ Glucose with 1uM Janelia Fluor^®^ 646 HaloTag^®^ ligand (Promega Corp.) and allowed to incubate at RT for 2 hours covered in foil. Following this, 0.5uL of the resuspension was then placed on a gel pad and the pad and prepared for imaging as specified in the “Live cell imaging” section below.

### Rapid Z-ring decondensation experiments

Cultures of strain RM077 and RM078 were started from a single colony and grown in LB at 37ºC overnight until the cells reached stationary phase. The overnight LB culture was then diluted 1:200 in 3mL of M9+ Glucose and grown overnight at room temperature (RT, 25ºC) until they reached log-phase (OD600 between 0.1 and 0.2) at which point the culture was induced with 10uM IPTG and 0.2% arabinose. Following a 2-hour induction at RT. Cells were prepared for imaging by centrifuging 1mL of the culture at 13k rpm for 1 minute and then resuspending the pellet in 100uL of fresh M9+ Glucose. Following resuspension, 0.5uL of the resuspension was transferred to a 3% M9+ Glucose agarose gel pad containing 10uM IPTG and 0.2% arabinose for imaging.

### Light induced inhibition of cytokinesis experiments

Cultures were started from a single colony of the strain of interest (see **Supplementary Table 4**) and grown in LB containing chloramphenicol and/or streptomycin (depending on the strain background, see **Supplementary Table 4**) at 37ºC overnight until the cells reached stationary phase. The overnight culture was then diluted 1:100 in fresh of M9+ Glucose containing chloramphenicol and streptomycin and was grown overnight at 25ºC until it reached log-phase (OD600 between 0.1 and 0.2) at which point the culture was induced. Following a 2-hour induction at 25ºC the cells were washed into fresh M9+ Glucose media lacking inducer and allowed to outgrow at 24ºC for 2 hours. After the outgrowth, the cells were prepared for imaging by centrifuging 1mL of the culture at 13k rpm for 1 minute and then resuspending the pellet in 40uL of fresh M9+ Glucose. Following this, 0.5uL of the resuspension was then placed on a gel pad and the pad and prepared for imaging as specified in the “Live cell imaging” section below. Control cells were placed in a chamber, wrapped in foil, left at RT in next to microscope for the duration of the experiment and imaged after the experiment ended.

### Live cell imaging

Live cell imaging was performed on a custom-built optical setup routed to an Olympus IX71 inverted microscope with a 100X 1.49 NA oil-immersion objective (Olympus Inc.). The light was focused onto the chip of an EMCCD camera (iXon Ultra 897, Andor Technology) with a final pixel size of 160 × 160 nm. The imaging focal plane was controlled by a piezo-driven stage (ASI, Eugene, OR). The EMCCD camera, lasers and shutters were controlled by Metamorph™ software (Molecular Devices).

Excitation light was provided by solid state 488nm (Coherent OBIS), 561 (Toptica Photonics) or 647nm (Coherent OBIS) lasers. The fluorescence emission signal was collected using either a ZET 488/561nm (Chroma Technology) or ZET 488/561/647nm (Chroma Technology) dual-band dichroic depending on the imaging conditions used. For two-color experiments, the emission light from the RFP and GFP channels were split using a 525/50 (Chroma Technology) and 650/50 (Chroma Technology) filter set mounted inside of an Optosplit II beam-splitter system (Cairn Research) prior to focusing on the EMCCD camera chip. The 488nm laser power used to activate the CRY2/CIBN system was the same as what was used to image GFP fusions 8.08 W/cm^2^ measured 1cm from the objective for PopZ and ZapA experiments and 80.8 W/cm^2^ measured 1cm from the objective for the TetR experiments measured using a Thor Labs Inc. Power Meter). The 561nm laser power used to image the mCherry fusions was 8.71 W/cm^2^ measured 1 cm from the objective for the PopZ and ZapA experiments and 93.3 W/cm^2^ measured 1 cm from the objective for the TetR experiments unless otherwise noted in the text.

For two-color experiments using the 647nm laser, the emission light from the RFP and GFP channels were split using a 556nm long-pass filter (T556lpxr, Chroma Technology) in the Optosplitter followed by 700/75 (Chroma Technology) filter for the RFP emission light prior to focusing on the EMCCD camera chip.

To avoid activation of the CRY2/CIBN system by the LED bright-field lamp (LDB100F System, Prior Scientific Inc.), we placed a 715nm long-pass filter (RG715, Thor Labs Inc.) or a 665nm long-pass filter (ET665lp, Chroma Technology) filter after the condenser and the BF lamp intensity was adjusted such that the measured 488 nm light after the filter was measured to be ∼200nW at 1cm from the sample plane.

Gel pads used to support the living bacteria cells were prepared by melting 3% w/v solution of agarose powder (SeaPlaque™ GTG™, Lonza Scientific) and either ERDM or M9+ Glucose for 0.5-1 hour at 70ºC. The melted gel was then transferred to a gasketed, cover-glass (FCS2, Bioptechs Inc.) and allowed to polymerize while sandwiched between the cover-glass and a cleaned coverslip (40 CIR-1, VWR Inc.) for 4 hours at 25ºC (room temperature). 0.5μL of the cells to be imaged were placed on the gel pad and allowed to dry (∼3 minutes) before a fresh coverslip was placed on top and the imaging chamber (FCS2, Bioptechs) was assembled.

For the DNA recruitment experiments, the cells were irradiated with 30 ms pulses of 488 nm and 561 nm light delivered consecutively every 5 seconds using a 30 ms exposure time for each.

For the cell pole and Z-ring recruitment experiments the cells were first imaged for 2.5 seconds with 561 nm light after which they were irradiated with 488nm light for 100ms and then imaged with 56 nm light by streaming for 15 seconds using a 50ms exposure time.

For the cell pole recruitment experiments where the green (561 nm) light was varied: the cells were first imaged for 2.5 seconds with both 647 nm and 561 nm light after which they were irradiated with 488 nm light for 100 ms and then imaged with 647 nm and 561 nm light by streaming for 15 seconds using a 50 ms exposure time.

For the fast Z-ring decondensation experiments, cells were initially imaged with 561nm light after which they were irradiated with 50ms pulses of 488nm light delivered consecutively every 10 seconds for a 5-minute period. The cells were then imaged again with 561nm light at the end of the 5-minute period.

For the Light-induced Inhibition of Cytokinesis (LInC) experiments, cells were irradiated with alternating 100ms pulses of 488nm and 561nm every five minutes for a period of 12 hours.

### Quantifying the RFP and/or GFP signal in cell pole and Z-ring recruitment experiments

A variation on a previously published custom MATLAB script^72^ used to quantify FRAP data was created to measure the GFP and RFP signals in the live-cell recruitment experiments. Briefly, cells were manually selected and cropped using a maximum intensity projection image of the entire recruitment stream (created in ImageJ) to avoid any selection bias. The max intensity image was also used to manually crop both the total cell area as well as the recruitment site area (either Z-ring or cell pole). For the Z-ring experiments the selected area was cropped manually cropped using a rectangular selection. For the cell pole experiments the PopZ region was cropped using a fixed circle with a 2-pixel radius.

We noticed that our experimental setup introduced a slight variation in the time between activation with blue light and acquisition with either green or far-red light. To accurately account for this delay, we created a custom MATLAB script that determined the true time delay between activation and acquisition frames using the imaging metadata obtained by Metamorph and then properly accounted for the delay when aggregating data and/or making measurements.

### Quantifying the percentage increase in the DNA recruitment experiments

To quantitatively characterize the accumulation of CRY2-mCherry at chromosomal sites, we calculated the increase in the Weber contrast^73^ of single cells. We noticed that the DNA spots were able to diffuse in both the lateral and azimuthal directions (relative to the *z*-axis of our objective). This type of movement made it difficult to track individual spots and accurately estimate their intensities using the method we applied to our cell pole and midcell platforms (see above). Since the accumulation of CRY2-mCherry from a uniform distribution to a single concentrated location inside the cell body is equivalent to an increase in the Weber Contrast of a single cell image, we estimated the accumulation using the following method. First, we segmented the cells with observable spots in the last frame of the time lapse movie. There are normally 1 or 2 spots in a segmented cell region. Then we subtracted the background intensity of each image using ImageJ^74^. Finally, the normalized contrast for a single cell, *j*, in frame *i* is defined as:

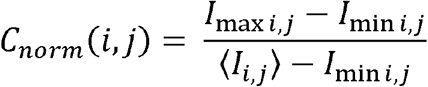

Where *C*_*norm*_(*i,j*) reflects the extended accumulation of CRY2-mCherry in the DNA spot and *I*_*max i,j*_ and *I*_*min i,j*_ are the intensity of the brightest and darkest pixels in the cells, respectively. The contrast is normalized by the mean intensity, ⟨*I*_*i,j*_⟩, of all the pixels in this region considering cell-to-cell variations in expression levels.

The enrichment curve is obtained by averaging all the cells in the same frame and dividing by the initial value to obtain the fold-increase. The value of 1 is subtracted from the fold-increase and that value is multiplied by 100 to obtain the percent increase. The error bars indicate the standard error of the mean. The curve is then fit by a single exponential function to calculate the average accumulation time.

### Quantifying the Halo signal in cell pole dissociation experiments

Individual well-isolated cells were cropped from whole field of view time-course image stacks starting 5 minutes after CRY2/CIBN association was activated using blue light. The individual cell image stacks were then registered in x-y space using the ImageJ StackReg^75^ plug-in to minimize the shifting of the cells over long timescales due to growth. The depletion of cell pole foci was then measured using the Weber contrast-based methodology employed for the TetR recruitment platform described above. The resulting traces for individual cells were averaged together and the exponential decay curve was fit to

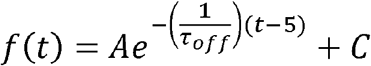

to extract the dissociation time constant (τ_off_).

### Quantification of Z-ring decondensation

Cells were segmented using the MiSiC deep learning-based cell segmentation algorithm^76^. After segmentation, the resulting cell masks were fed into MicrobeJ^77^ to quantify the mean ZapA-mCherry intensity at across the long axis of the cell and cell lengths. All cells were visually inspected to remove any cells that lacked a Z-ring at the start of the experiment. The midcell localization fraction of the Z-ring was measured by dividing the integrated ZapA-mCherry fluorescence intensity within a 3-pixel window (480 nm width) about the midcell maximum (approximate Z-ring position) by the total integrated ZapA-mCherry fluorescence intensity of the whole cell both before and after a 5-minute blue light induction. The change in the midcell localization fraction before and after blue light induction was calculated, normalized by the midcell fraction before induction and multiplied by a factor of 100 to obtain the percent change in Z-ring intensity at midcell. The percentage of cells that both harbored a Z-ring prior to induction with blue light and showed a reduction in the midcell fraction of ZapA-mCherry intensity at midcell greater than 4% were deemed as “decondensed”. This decondensation threshold was applied to every cell in each experimental population to determine the fraction of cell that underwent significant decondensation in each experimental condition. Bootstrapping was employed to estimate the standard error of the mean. Briefly, a subpopulation of cells (500 for the induced cells and 300 for the uninduced cells) were randomly sampled and used to calculate the percentage of cells that underwent decondensation. This calculation was repeated 100 times to obtain the standard error of the mean.

### Quantification of inhibition of cytokinesis

Cells were manually tracked throughout the duration of the experiment to determine if and when a successful division occurred. Cells were cropped and the average midcell intensity across the long-axis of the cells (long-axis projection) was calculated using a custom MATLAB script.

### Quantification of maximal enrichment and reduction of CRY2-mChery and CRY2-HaloTag

In order to reduce the effects of noise on the maximal fold enrichment and maximal fold reduction of CRY2-HaloTag signal at the cell poles (*i*.*e*., estimation of the plateau value), the final plateau value was calculated by taking the mean of the last ten datapoints of individual cell trajectories.

## Supporting information

Supplementary Video 8 (GFP Channel)

Supplementary Video 8 (RFP Channel)

## SUPPLEMENTARY MATERIAL

## ACKNOWLEDGMENTS

We would like to thank T. Bernhardt and J. Buss for the BW25113 (*zapA-mCherry*) strain. We would also like to thank E. D. Goley for providing the *Cc*PopZ, *Cc*FtsZCTL, *Atu*FtsZCTL sequences and for providing valuable input during discussions about this work. Additionally, we would like to thank N. Yehya for helpful insight and discussions. This work was supported by NIH R01 GM086447, NIH R35 GM136436 (to J.X.) and NIH T32 GM008403 to (R.M.).

## AUTHOR CONTRIBUTIONS

R.M. constructed and characterized all the strains and fusions and proposed and carried out all experiments as well as quantified and interpreted all data unless otherwise noted. C.H.B. assisted in developing methodologies for fold enrichment calculations with R.M. and J.X and assisted in analysis of CRY2/CIBN green light effects. X.Y. developed the Normalized Contrast method used to analyze the DNA localization data and performed the data analysis using this method. J.W.M., together with R.M., designed strains for the light induced inhibition of cytokinesis assay. J.X. planned and directed the project and, together with R.M., designed experiments, interpreted the data and wrote the manuscript.

## COMPETING FINANCIAL INTERESTS

The authors declare no competing financial interests.

## Notes

### Competing Interest Statement

The authors have declared no competing interest.

