## Supplementary Figure 1 for "Light-dependent modulation of protein localization and function in living bacteria cells"

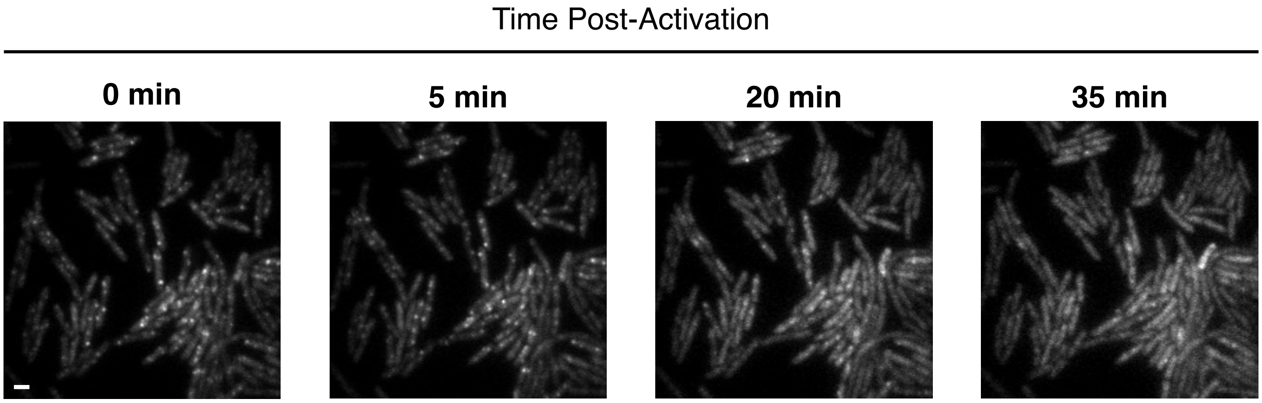


**Supplementary Figure 1 I Relaxation of the CRY2/CIBN system in the absence of blue light.** The CRY2/CIBN system in IL02 cells exogenously expressing TetR-CIBN and CRY2-mCherry was fully activated with blue light ($t$ = 0) demonstrated by the formation of DNA foci within the cell body and allowed to relax in the dark while being imaged every 5 minutes with green (561nm) light to visualize the distribution of CRY2-mCherry. After ~35 min in the dark CRY2-mCherry fluorescence had relaxed back to a uniform distribution within the cell body. Scale bar = 2µm.

### Supplementary Figure 1
