## Supplementary Figure 2 for "Light-dependent modulation of protein localization and function in living bacteria cells"

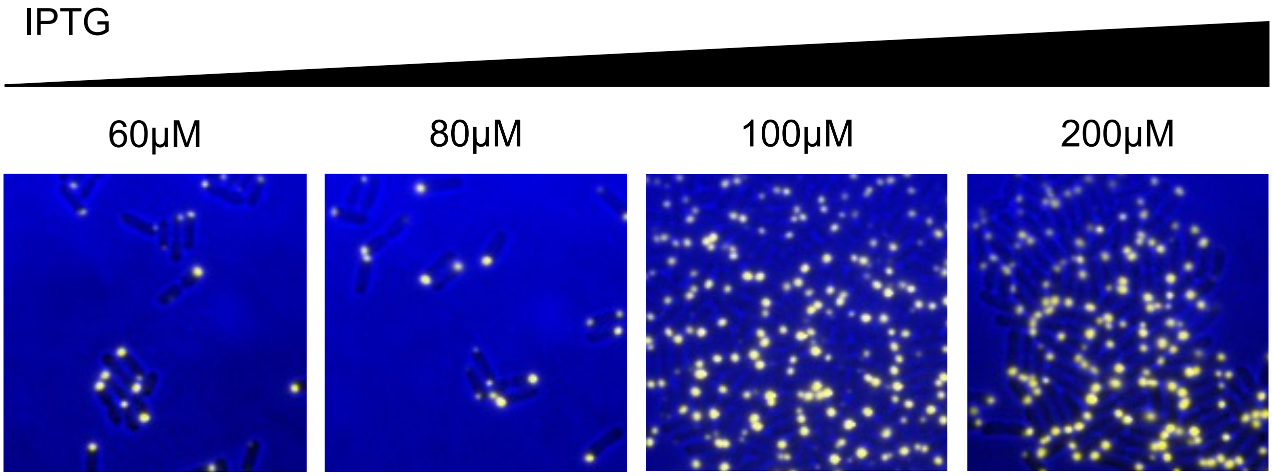


**Supplementary Figure 2 | CIBN-GFP-PopZ foci formation at different expression levels.** Cells (gray) exogenously expressing CIBN-GFP-PopZ (yellow) from a *lac*-inducible promoter were induced under different IPTG concentrations for an hour at 37°C. As the concentration of IPTG increased, larger and more stable CIBN-GFP-PopZ foci began to form more homogenously over the cell population. Here, we chose 100μM IPTG as the optimal induction level to maintain a high level of PopZ foci formation across the cell population.

### Supplementary Figure 2
