## Supplementary Figure 3 for "Light-dependent modulation of protein localization and function in living bacteria cells"

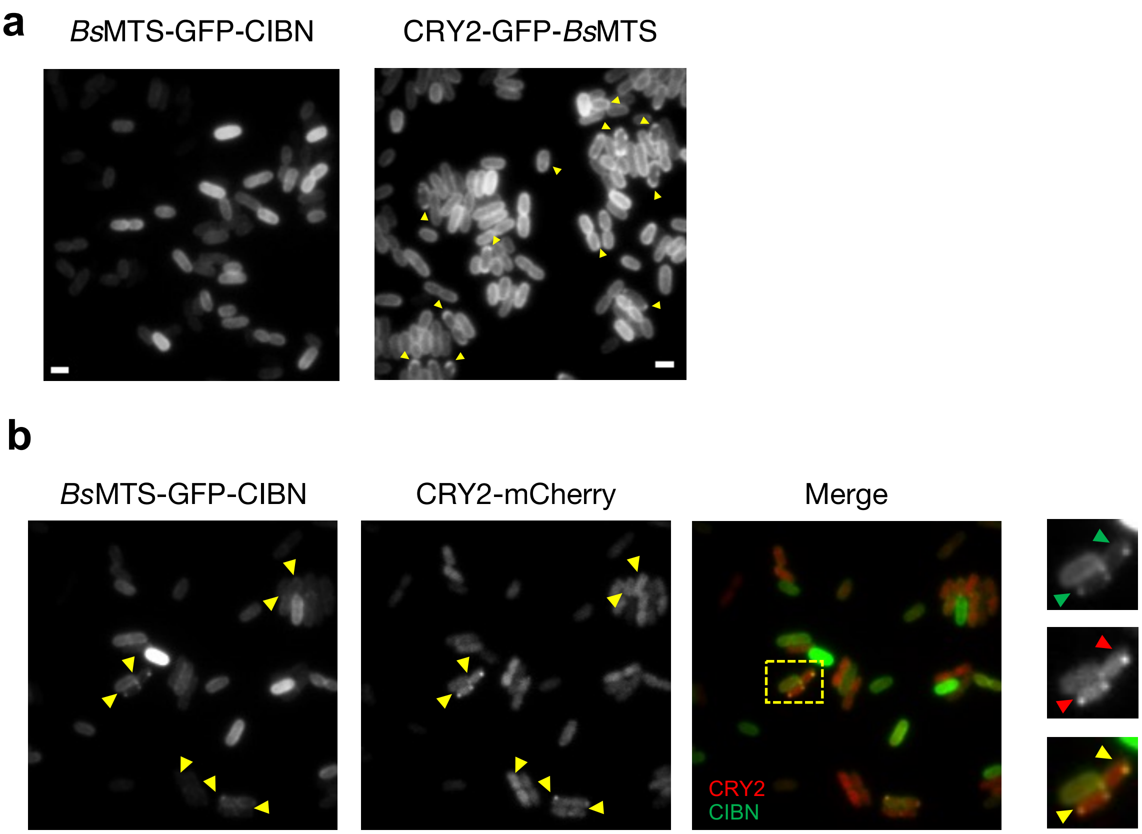


**Supplementary Figure 3 l Recruitment of cytoplasmic protein cargo to the inner membrane (M).** (**a**), Example images of cells exogenously expressing a *Bs*MTS-GFP-CIBN fusion from a *lac* promoter (*left*) and CRY2-GFP-NsMTS from an *araBAD* promoter (*right*). Both membrane-targeted proteins show the expected uniform membrane distribution. (**b**), IM-associated puncta formation of *Bs*MTS-GFP-CIBN and CRY2-mCherry after blue light activation. Yellow arrowheads point to colocalized BsMTS-GFP-CIBN and CRY2-mCherry puncta in both color channels (Merge). Rightmost column: Zoom-ins of example cells (boxed in Merge) exhibiting CRY2/CIBN co-localization. Scale bar = 2µm.

### Supplementary Figure 3
