## Supplementary Figure 4 for "Light-dependent modulation of protein localization and function in living bacteria cells"

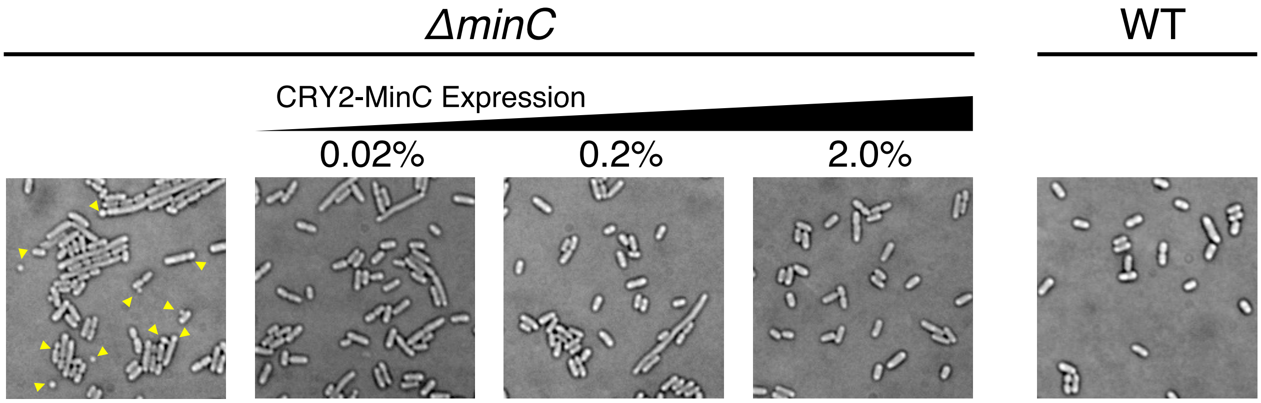


**Supplementary Figure 4 l CRY2-MinC Functionality Test.** A *minC* null strain without exogenous expression of CRY2-MinC fusion shows the canonical minicell and polar constriction phenotypes (yellow arrows). These phenotypes were rescued and cells exhibited wildtype (WT)-like morphology when CRY2-MinC was exogenously expressed from an arabinose-inducible promoter even at a low 0.02% arabinose induction level.

### Supplementary Figure 4
