## Supplementary Figure 5 for "Light-dependent modulation of protein localization and function in living bacteria cells"

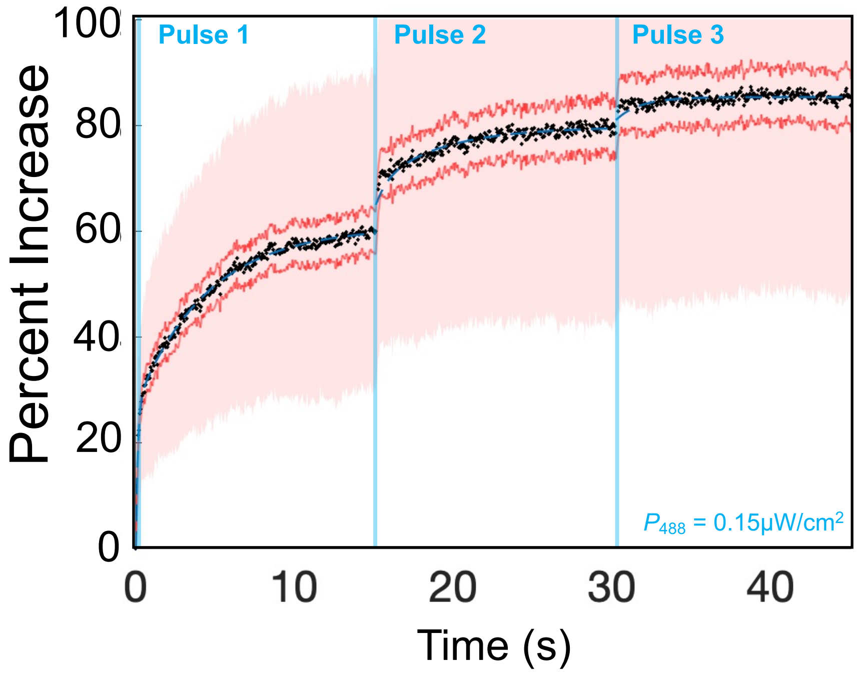


**Supplementary Figure 5 | Pulse induction of CRY2/CIBN association using variable blue light conditions.** Mean percentage increase of CRY2-HaloTag(JF646) fluorescence at the cell pole from individual cells where CRY2/CIBN association was stepwise activated using multiple 100 ms blue light pulses of the same power. Solid lines indicate the s.e.m. whereas the transparent overlay indicates the s.d.

### Supplementary Figure 5
