## Supplementary Figure 6 for "Light-dependent modulation of protein localization and function in living bacteria cells"

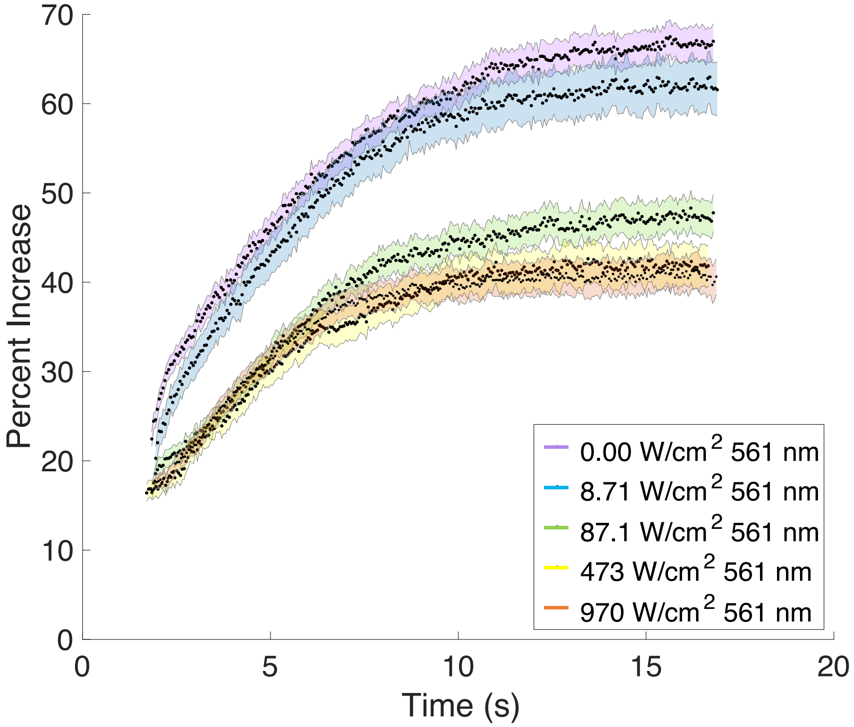


**Supplementary Figure 6 | Effect of Green Light on CRY2-Halo Polar Enrichment.** Averaged percent increase traces of cell pole-recruited CRY2-Halo under varying green light intensities shows a green light dependent reduction in the final plateau but not the $\tau_{90}$ time constant (error bars = s.e.m. **Supplementary Table 3**).

### Supplementary Figure 6
