## Supplementary Figure 7 for "Light-dependent modulation of protein localization and function in living bacteria cells"

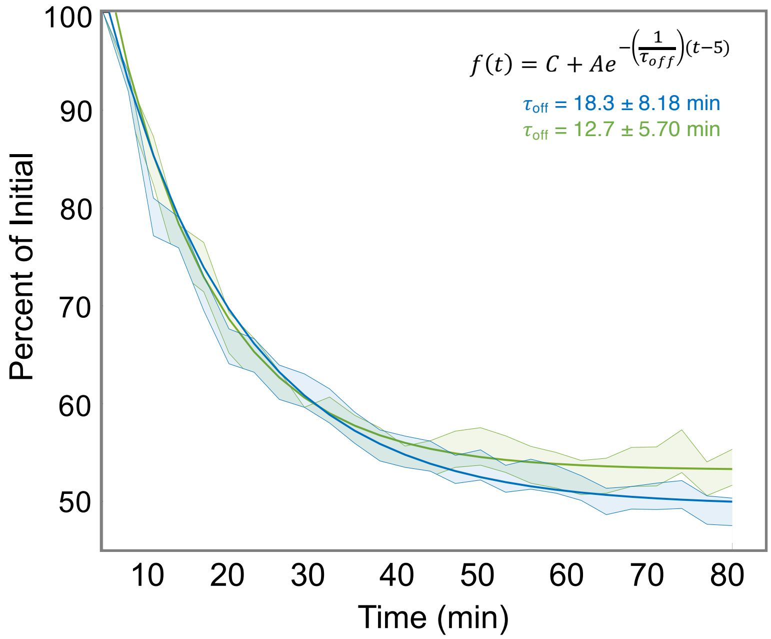


**Supplementary Figure 7 I The effect of green light on CRY2/CIBN dissociation.** Quantification of the percentage of CRY2-HaloTag at the cell poles harboring a CIBN-GFP-PopZ fusion after full induction by a 100ms activation pulse of blue light. A snapshot was taken every 3 minutes post-induction using either 647 nm light only (blue curve, $\tau_{off}$ = 18.3 s ± 8.18 min) or 674 nm light delivered with 561 nm light (green curve, $\tau_{off}$ = 12.7 ± 5.70 s,). The dissociation curves were fit with a single exponential decay to obtain the corresponding time constants. Shaded regions indicate s.e.m.

### Supplementary Figure 7
