## Supplementary Note 1 for "Light-dependent modulation of protein localization and function in living bacteria cells"

For the purpose of the LInC assay, expression levels of CRY2-MinC and ZapA-CIBN in cells need to be optimized so that cell division is not inhibited until light induced complex formation between CRY2 and CIBN brings MinC to the Z-ring. On one hand, a very high level of overexpression of MinC is known to destabilize of WT Z-rings by itself, resulting in cell filamentation and lysis^1^. On the other hand, overexpression of ZapA is known to over-stabilize the Z-ring that likely rendering it more resistant to MinC’s activity^2^. Therefore, the expression levels of both of ZapA and MinC fusions need to be carefully controlled so that the Z-ring morphology and cell growth/division are not perturbed prior to light induction, but efficiently perturbed after light induction.

To achieve optimal expression levels of both CRY2-MinC and ZapA-CIBN, we grew cells harboring ZapA and MinC fusions overnight in LB to reach saturation. We then diluted the overnight culture 1:200 in M9+ medium and allowed them to reach early log-phase (OD_600_ between 0.1 and 0.2). We then shifted the culture to room temperature to slow down their growth and induced the culture with different concentrations of IPTG and arabinose for 1-2hrs. After the induction period we immediately examined the cell and Z-ring morphology in the absence of blue light. From these experiments, we determined that the induction condition of 10μM IPTG for ZapA-CIBN and 0.2% arabinose for CRY2-MinC yielded the most WT-looking cells as determined by cell length measurement.
