## Supplementary Table 1 for "Light-dependent modulation of protein localization and function in living bacteria cells"

**Supplementary Table 1 | Overview of inner membrane targeting methods explored.** Briefly, different membrane targeting sequences (ZipA, Tsr or FtsN), CIBN or CRY2 fusion, N or C-terminal fusion, expression levels and light activation conditions were tested. See reference 6 for more detail.

| CIBN Fusion | CRY2 Fusion | Fusion Description | Experimental Result Summary |
|---|---|---|---|
| ZipAmts | mCherry | ZipA(1-183); early division protein, transmembrane domain with a flexible linker[1]. | No CRY2/CIBN co-localization observed. |
| Tsr | mCherry | Chemotaxis receptor, transmembrane protein[2]. | Dynamic polar puncta observed. |
| mCherry | FtsN | Late division protein, transmembrane protein[3,4]. | No CRY2/CIBN co-localization observed. |
| FtsNtmd | mCherry | FtsN(34-53); late division protein fragment, transmembrane domain[4]. | No CRY2/CIBN co-localization observed. |
| n/a | CRY2olig-mCherry CRY2olig-FtsNtmd | FtsN(34-53); late division protein fragment, transmembrane domain, CRY2oig is a self-clustering mutant of CRY2[5]. | No CRY2/CIBN co-localization observed. |


### Supplementary Table 1
