## Supplementary Table 2 for "Light-dependent modulation of protein localization and function in living bacteria cells"

| | Cell Length (um) | 1.51 | 2.55 | 3.58 | 4.61 | 5.65 | 6.68 | 7.71 | 8.75 | 9.78 | 10.82 |
|---|---|---|---|---|---|---|---|---|---|---|---|
| **Activation Light On** | **% Change Z-ring Int.** | 0.93 | -0.89 | 0.41 | -2.37 | -4.69 | -12.64 | -7.27 | -1.52 | -11.45 | -8.47 |
| | **SEM** | 0.58 | 0.67 | 0.88 | 1.13 | 2.06 | 3.95 | 6.36 | 7.38 | 9.36 | 2.17 |
| | ***N* cells** | 418 | 513 | 372 | 207 | 85 | 35 | 17 | 7 | 3 | 33 |
| **Activation Light Off** | **% Change Z-ring Int.** | 0.60 | 0.57 | 2.05 | 2.28 | 4.32 | -6.67 | -3.92 | -10.06 | -13.76 | -13.76 |
| | **SEM** | 0.55 | 0.61 | 0.92 | 1.39 | 2.88 | 3.67 | 4.27 | 4.23 | 5.00 | 1.56 |
| | ***N* cells** | 438 | 457 | 313 | 149 | 59 | 24 | 15 | 6 | 4 | 32 |

**Supplementary Table 2 | Cell Length Dependence of Short Timescale LInC System Activity.** Values of the percent reduction of ZapA-mCherry intensity (μ ± s.e.m.) at midcell of cells expressing the LInC system after either a 5 minute exposure of blue activation light (blue) or darkness (grey) as a function of cell length (see **Fig. 3h**).

### Supplementary Table 2
