## Supplementary Table 3 for "Light-dependent modulation of protein localization and function in living bacteria cells"

| | Green Light Intensity (W/cm²) | 0 | 8.71 | 87.1 | 437 | 970 |
|---|---|---|---|---|---|---|
| Association | % Increase Plateau | 68 | 59 | 46 | 42 | 41 |
| | SD | 10 | 10 | 5 | 5 | 1 |
| | Fold Reduction in Plateau Relative to No Green Light | 1 | 0.87 | 0.67 | 0.61 | 0.60 |
| | SD | 0.21 | 0.20 | 0.13 | 0.12 | 0.09 |
| | $\tau_{0.9}$ (s) | 7.63 | 6.91 | 6.59 | 6.40 | 6.46 |
| | SD | 0.64 | 1.53 | 1.58 | 1.31 | 0.16 |
| | Fold Reduction in $\tau_{0.9}$ Relative to No Green Light | 1 | 0.90 | 0.86 | 0.84 | 0.85 |
| | SD | 0.12 | 0.24 | 0.24 | 0.21 | 0.13 |
| | *N* cells | 286 | 184 | 170 | 168 | 182 |
| Dissociation | % Decrease Plateau | 51 | | | | 53 |
| | SEM | 1.8 | | | | 7.6 |
| | Fold Reduction in Plateau Relative to No Green Light | 1 | | | | 1.04 |
| | SEM | 0.05 | | | | 0.10 |
| | $\tau_{off}$ (min) | 18.3 | | | | 12.7 |
| | SEM | 8.18 | | | | 5.7 |
| | Fold Reduction in $\tau_{off}$ Relative to No Green Light | 1 | | | | 0.69 |
| | SEM | 0.34 | | | | 0.19 |
| | *N* cells | 91 | | | | 62 |

**Supplementary Table 3 | Green Light Effects on CRY2/CIBN Association and Dissociation.** (*Top*) Calculated values of the maximal percent increase at the cell pole and 90% recruitment time ($\tau_{0.9}$) of CRY2-Halo at varying green light intensities after blue light activation. (*Bottom*) Calculated values of the maximal percent decrease at the cell pole and apparent off time constant ($\tau_{off}$) of CRY2-Halo during the dissociation process of the pre-formed complex in the absence of presence of green light.

### Supplementary Table 3
